# Diffusion Weighted Image Co-registration: Investigation of Best Practices

**DOI:** 10.1101/864108

**Authors:** David Qixiang Chen, Flavio Dell’Acqua, Ariel Rokem, Eleftherios Garyfallidis, David J. Hayes, Jidan Zhong, Mojgan Hodaie

## Abstract

The registration or alignment of diffusion weighted images (DWI) with other imaging modalities is a critical step in neuroimaging analysis. Within-subject T1-DWI co-registration is particularly instrumental. DWI-derived scalar images are commonly used as intermediates for T1-DWI co-registration, and the resulting registration transforms are applied to all other scalar images for analysis. The ideal registration intermediate should register well to T1 and other multimodal images and be practically easy to obtain. It is however, currently unclear which DWI-derived scalar image serves as the best intermediate. We aim to determine the best, practical, intermediate for image co-registration. T1 and DWI images were acquired from 20 healthy subjects. DWIs were acquired with 60 directions. Six DWI-derived scalar images were compared including: 1) fractional anisotropy (FA); 2) generalized FA (GFA); 3) B0 images; 4) mean DWIs with the B0 image (MDWI); 5) anisotropic power (AP) images. AP showed the smallest variability in registration improvements across all the tested DWI derived scalar images, and show the highest average percent changes with CC registration cost function (CC=1.2%, MI=15%). In contrast, the FA and GFA transforms resulted in significantly poorer registration across DWI types. The AP image was the DWI-derived scalar image that provided the most consistent registration to all other images. Practically, it is generated easily and so could be implemented in basic and clinical research pipelines currently using other intermediates. Given these findings, it is recommended that AP images be used for T1–DWI co-registration, and that FA and GFA images in particular be avoided.

## 2. Introduction

Diffusion magnetic resonance imaging (dMRI) allows for visualization and quantification of the brain’s white matter by measuring the anisotropy of water molecules (Lenglet et al., 2009). The resulting diffusivity parameters are used to infer white matter directionality which are visualized using diffusion tensor images (Basser & Jones, 2002). These form the basis of study for a wide range of disease processes (Chen et al., 2016a; Ciccarelli et al., 2008; Hodaie et al., 2012). DMRI has become an essential tool in the in vivo analysis of brain white matter, particularly when used at the group level (Chen et al., 2016b) to compare diffusion metrics and to model white matter tracts using tractography (Chen et al., 2016b, 2015; Hodaie et al., 2010). An increasing number of neuroimaging techniques, including structural connectivity analysis, also depend on dMRI to segment white matter regions (Moayedi & Davis, 2012; McGrath et al., 2013; Wiech et al., 2014).

Image registration permits the transformation of different, individual, diffusion images into one brain template and allows for group analysis. This multi-step process involves the accurate co-registration of within-subject diffusion-weighted images (DWI) and anatomical T1-weighted (T1) images, followed by registration of between-subject T1s to create a common template. Individual T1 images typically serve as the intermediate space due to their high spatial resolution and low incidence of distortions (Avants et al., 2009; Brown, 1992; Klein et al., 2009; Tustison et al., 2014). The registration of T1s to DWIs is critical to ensure the validity, reliability, and interpretability of the final results (e.g., in correlating DWI data to other measures, and for use in group analyses e.g. (Chen et al., 2016b). However, while T1 to DWI co-registration is a common procedure in neuroimaging studies, it is currently unclear which DWI-derived scalar image serves as the best intermediate.

The greatest challenge of T1-DWI co-registration is that DWI acquisitions are susceptible to both affine/linear (i.e. eddy-current and head motion) and non-linear echo planar image field distortions (Rohde et al., 2004). The most common strategies to account for such issues include: 1) correcting all DWI distortions before co-registration with a T1 image, and/or 2) using non-linear co-registration transformations to best warp the anatomical image to DWI space. Corrections for DWI distortions generally involve affine registration of each of the diffusion gradient images to a non-diffusion-weighted image (B0), followed by a rotational correction of the original diffusion gradient b-matrix (Leemans & Jones, 2009). This approach, however, does not account for non-linear distortions, such as those typically found in the brainstem and frontal and temporal cortices, that are due to MRI field inhomogeneity. Affine co-registration of the T1 image directly to DWI results in poor and highly variable overlaps – increasing the likelihood of both type I and type II errors.

One strategy is to calculate an “anti-distortion” image by acquiring either an extra set of B0 images, or a full DWI sequence with reversed phase-encoding, followed by the construction of a displacement field map which provides an estimate of the undistorted DWI using a least squares (Andersson et al., 2003), diffeomorphic (Irfanoglu et al., 2015), or Gaussian approach for acquisitions with high b-values (Andersson & Sotiropoulos, 2015). However, most prevailing MR datasets, particularly in the clinical domain, still use lower b-values, and lack reversed B0 acquisitions. In such cases, the most commonly used strategy to minimize errors is to nonlinearly co-register the T1 image to an intermediate DWI-derived scalar image. Although it is also possible to indirectly perform T1-DWI co-registration through high resolution T2-weighted images, as with the other issues, clinical researchers rarely acquire such scans. As such, we have focused exclusively on direct within-subject DWI-T1 co-registration solutions. An additional important consideration is the ability of a registration transform to improve the registration for all relevant DWI-derived scalar images. In addition to the distortion issue, a reliable intermediate image must also be chosen for T1-DWI co-registration, from which the registration transform can be derived. Importantly, since MR image intensities can be inverse (e.g. T1-like and T2-like intensities), registration improvements may not be uniform, and instead vary depending on the intensity profiles similarities. Despite these concerns, the co-registration utility of the most common, easily producible, DWI-derived intermediates has not yet been investigated.

### 2.1. Image Intermediates

Currently, the five most common DWI-derived scalar images, which are most likely to be of practical use as co-registration intermediates, include: 1) fractional anisotropy (FA) images (Basser & Jones, 2002; Sboto-Frankenstein et al., 2013); 2) generalized FA (GFA) images; 3) non-diffusion-weighted B0 images (B0); 4) mean DWIs (MDWI) where the DWI acquisition, often represented as a 4D image, is averaged across the sequences; 5) and the anisotropic power (AP) image (Dell’Acqua et al., 2014), which is an anisotropy map derived from the spherical harmonics estimate coefficients of high angular resolution DWI images. AP directly derives a T1-like contrast image from the diffusion weighted image itself, and therefore shows promise in improving T1-DWI co-registration. Some images, such as B0 and mean DWI, are derived directly from the DWI acquisition volume, while others (FA, GFA, AP) are parametric image maps indirectly calculated from mathematical diffusion models.

The simplest scalar image is the B0 images as part of the DWI acquisition. B0 is used as part of the pipeline for grouped diffusion studies (Gupta et al., 2016; Yeatman et al., 2012)and to register T1 anatomical to diffusion tensor images for structural connectivity studies (Cao et al., 2013). Since B0’s intensity profile matches well with T2 anatomical images, there also has been attempts to use inverse intensity to optimize registration with T1 using B0 (Bhushan et al., 2015).

The second set of scalar images is derived by taking the mean of the DWI dataset (MDWI) by collapsing the gradient dimension, which transforms the 4D dataset to 3D. MDWI is commonly used in multi-modal registration pipelines (Peng et al., 2009).

Fractional anisotropy (FA) image (Basser & Pierpaoli, 2011) is also commonly used as a registration intermediate. Due to its high intensity in white matter regions, FA is commonly used with probabilistic tractography registration to standard T1 space (Mansour et al., 2013; Salomons et al., 2012). Research also shows that normalized cross-correlation and template matching can improve FA-T1 registration (Malinsky et al., 2013).

An alternative to FA is to use the generalized fractional anisotropy (GFA) image. Since FA shows low intensity in cross-fibre regions, it may adversely affect registration in those areas. GFA can properly characterize crossing-fibre intensities, and therefore may be a better candidate than FA. The GFA is computed from the SH coefficients of spherical harmonics based methods such as Q-ball, where the diffusion orientation distribution function (ODF) is denoted as Ψ(*u*) (Tuch, 2004), such that:

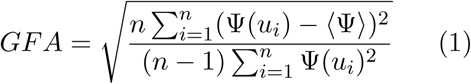

where n is the number of gradient directions.

Another promising candidate is the anisotropic power (AP) image (Dell’Acqua et al., 2014). AP have similar visual intensity to T_1_, and can be computed from HARDI acquisitions without any acquisition modifications. Since AP are derived from the SH model, it is also capable of proper cross-fibre region characterization. AP images are derived from the even harmonic orders (l) of SH coefficients (Descoteaux et al., 2006; Frank, 2002) and are defined as

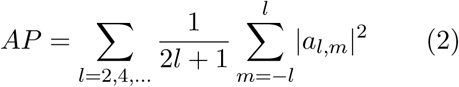

where l is the harmonic order, m is the l-th order coefficient index, a is the corresponding SH coefficient. The AP values are then normalized to non-negative scale by the function:

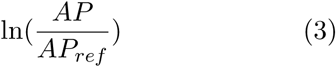

where AP_ref_ is the normalization constant. The resulting image will closely match T_1_ intensities, and can better capture contrasts in the gray matter structures. AP values stabilize at 6 harmonic orders (28 directions), meaning that AP images can be reliably derived from HARDI sequences with more than or equal to 28 gradient directions.

### 2.2. Quantification of Image Registration

Image registration methodologies involve iterative processing aimed at minimizing the output of a cost function reflecting overlying image differences, thus maximizing the similarity between reference and moving images. Two commonly used cost functions for assessing similarity in medical imaging are cross-correlation (CC) and mutual information (MI) (Liu et al., 2013; Mercier et al., 2012). Mutual information is based on the Shannon entropy score, measuring the dispersion of values between two images (Pluim et al., 2003), and by extension represents the likelihood that two images share mutually dependent information (Mattes et al., 2003). Cross-correlation measures image similarities by the profile of their intensities (Maintz & Viergever, 1998). While CC appears to be a more sensitive measure of matching local features, MI is thought to be a more global measure of similarity (Crum, 2004). Furthermore, MI may be the optimal cost function for affine registration, while CC may be a better choice for non-linear registration where local feature-matching is of greater importance (Avants et al., 2011). Unlike T1 anatomical registration comparisons (Klein et al., 2009), where automated segmentation tools are available to serve as an independent metric of comparison, there is currently no automatic method available in DWI-T1 space that can be used to judge the quality of registration. As the best cost function to use for T1-DWI co-registration remain unclear, we use both CC and MI to independently assess the registrations.

There is currently no consensus in the literature about which intermediate image should be used for T1-DWI image registration purposes. This information is pertinent to ensure the increased validity, reliability, and interpretability of neuroimaging results. Our present study aims to quantitatively evaluate the use of each readily available DWI-derived scalar image type (i.e. B0, MDWI, FA, GFA and AP) in the registration process, using CC and MI cost functions as quantitative assessors, in order to determine the most efficient image type for registration and its ability to translate registration improvements to other image types.

## 3. Methods

### 3.1. Acquisition and Preprocessing

T_1_ and DWIs were acquired from 20 healthy subjects (mean age 31.1 ± 10.2; 10 females). Ethics approval was granted by the University Health Network Research Ethics Board (Toronto, Canada), MR images were acquired at the Toronto Western Hospital, and all subjects gave their informed written consent. DWIs were acquired on a GE HDx 3 Tesla MRI scanner, 8-channels head coil, with 60 directions and 1 non-diffusion-weighted acquisition using the following parameters: 0.9375×0.9375×3 mm3 resolution, matrix=256×256, b=1000 s/mm^2^, field of view (FOV)=240mm, TE=86.4 ms, TR=17000 ms, flip angle=90 deg. T_1_s were acquired with 0.9375×0.9375×1 mm^3^ resolution, slice spacing=1 mm, TE=5.052 ms, TR=11.956 ms, flip angle=20 deg, FOV=240 mm, and matrix=256×256. DWIs were initially corrected for eddy-current and head motion with affine registration using FSL FLIRT (Jenkinson & Smith, 2001), and appropriate rotational corrections to gradient vectors (Leemans & Jones, 2009) using in house software written in Python Numpy.

### 3.2. DWI Derived Scalar Image Processing

Brain masks were created for T_1_ and DWI using FSL bet (Smith, 2002), with bet fraction=0.2. Brain masks were contracted by a 2 mm spherical kernel in FSL to minimize the high-intensity skull halo that occurs in image types such as FA that may interfere with registration. The brain mask is then applied to all DWI derived scalar images.

B0, ADWI, MDWI, FA, GFA and AP images were each generated (Figures 1, 2) as follows: B0, ADWI and MDWI images were created using FSL (Smith et al., 2004); FA images were created in 3D Slicer by calculating the scalars following diffusion tensor estimation (Pieper et al., 2006); GFA and AP images were created with Dipy software library (Descoteaux et al., 2007; Garyfallidis et al., 2014).

**Figure 1:**
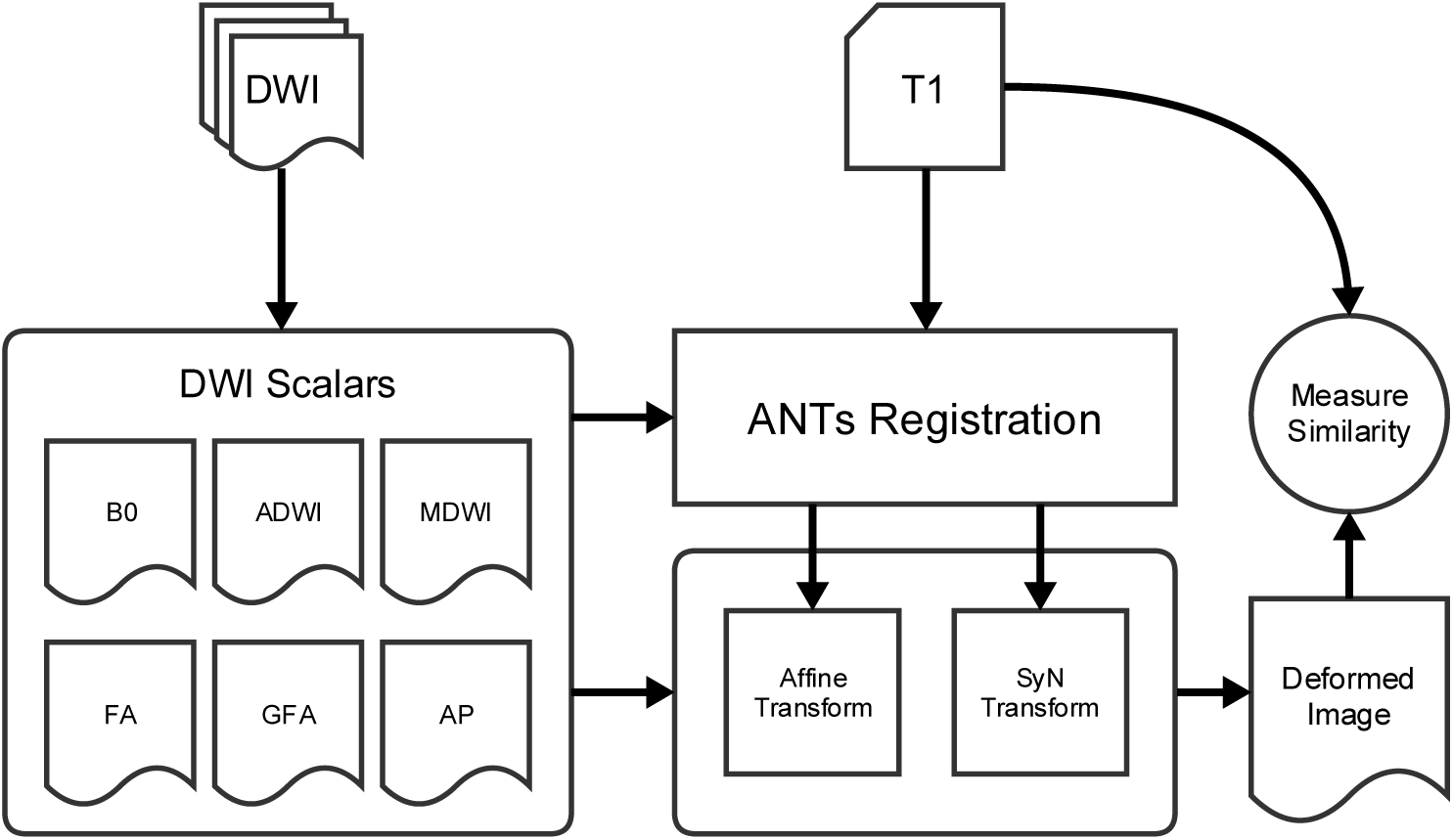
Illustrated overview of the processing steps to measure DWI to T1 registration similarities.

**Figure 2:**
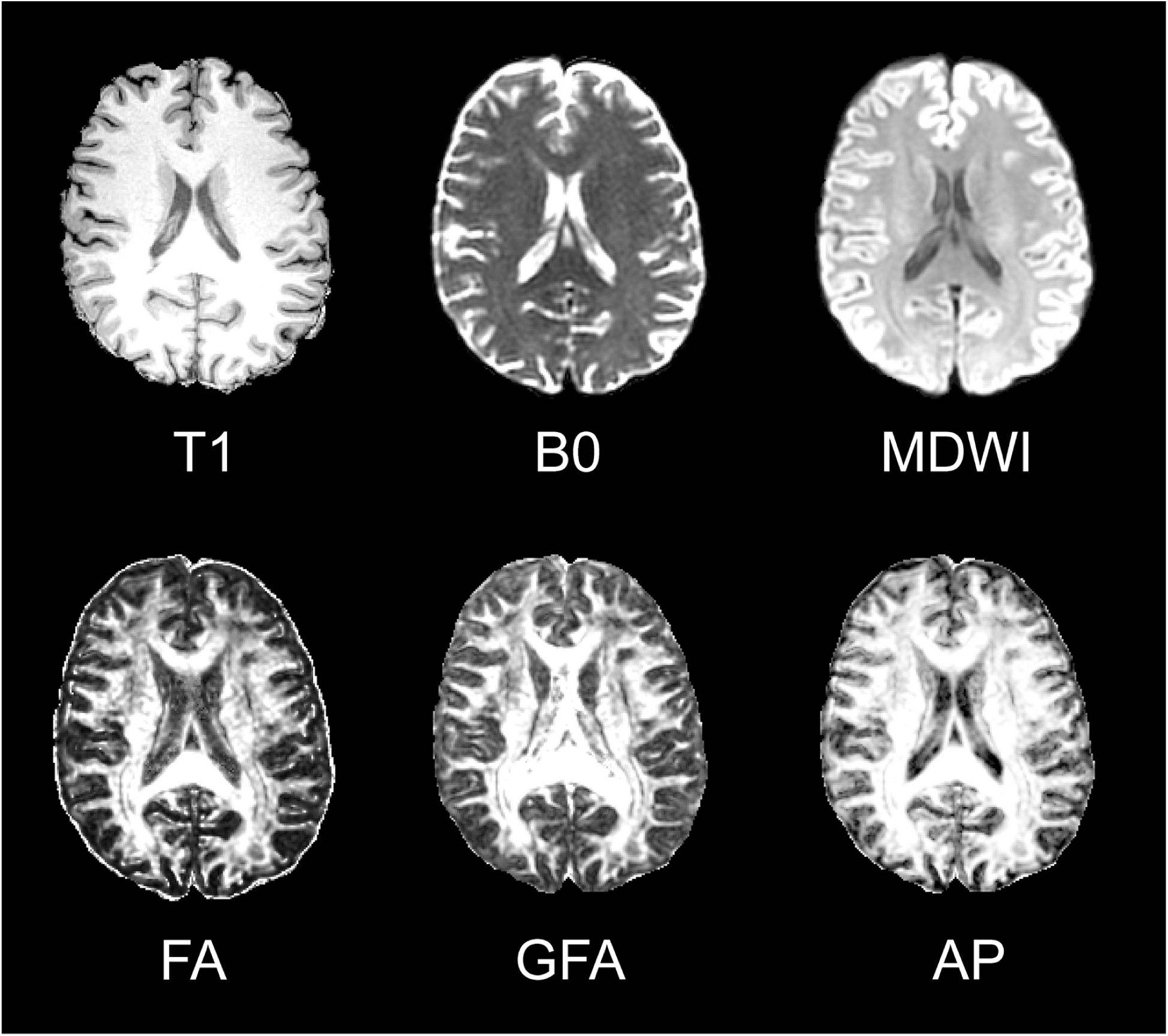
DWI derived scalar images (B0, MDWI, FA, GFA, and AP) comparing to T1 (top left).

Two sets of AP images were generated with an author-implemented AP algorithm in Dipy. The first (denoted as AP1) is created with matching normalization constant with the original method from Dell’Acqua et al (Dell’Acqua et al., 2014) (AP_ref_ = 10-5, then zero all image values <0). The second (denoted as AP2), is created with AP_ref_ = 1, and then the entire image values are shifted to the positive range based on the image minimum. Pair-wise log joint histogram of the DWI derived scalar images was plotted (Hunter, 2007) to evaluate image intensity orthogonality.

### 3.3. T1 to DWI Co-registration

Registrations were performed using Anatomical Normalization Tools (ANTs) (Avants et al., 2008) with affine and symmetric diffeomorphic registration. MI was used as the cost function for affine registration, while CC and MI were used separately as the SyN registration cost-functions. Parameters for affine registration: step size = 0.1, metric = mutual-information (MI), convergence = 10000 × 10000 × 10000 × 10000 × 10000, shrink factors = 5 × 4 × 3 × 2 × 1, smoothing sigmas = 4 × 3 × 2 × 1 × 0 mm. Parameters for symmetric diffeomorphic registration (SyN): MI metric weight = 1, MI bins = 32; CC metric weight = 1, CC radius = 3; convergence = 50 × 35 × 15,1e-7, shrink-factors = 3 × 2 × 1, smoothing sigmas = 2 × 1 × 0 mm, and use-histogram-matching = true.

The resulting registration transforms from each DWI derived scalar image were applied to all DWI derived scalar image types including themselves using 3rd order B-spline interpolation, where they were projected into T_1_ space. For each DWI scalar image, an identity transformation (*T*_identity_), affine-only transform (T_Affine_), and affine with SyN transforms (T_Affine_+SyN) were applied separately. MI and CC similarity metrics between the transformed DWI scalar and T_1_ images were obtained using ANTs. Percent change in both CC and MI similarity scores for each type of transformed images were calculated as:

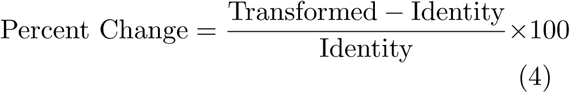

The higher the percent change is, the more similar the transformed DWI scalar and T_1_ images are after accounting for the identity transformation. ANOVA and Tukey post-hoc tests were performed using R statistics software (R Core Team, 2014) to compare differences in percent changes between similarity scores of images deformed by their own T_Affine_+SyN (autodeformation: D_auto_; e.g. comparing pre-to post-FA scores when FA image was used as the intermediate) as well as similarity scores across groups (D_all_; e.g. comparing scores resulting from using the FA intermediate to transform all other image types).

Additionally, we used a joint log-histogram of all the DWI derived scalar images to examine possible group biases in DWI scalar types. We intuited that the scalar maps can roughly be divided into two primary groups: those that are directly computed from the DWI directions (DSG), and share similar intensity profiles, such as B0, MDWI, ADWI; and those computed from anisotropic diffusion models (ASG), such as FA, GFA, and AP. It’s possible that there exists a degree of orthogonality between image types, such that there may be no single image type that can equally translate the registration improvements to other images with different intensity profiles. The joint log-histogram is to check for the existence of this type of orthogonality.

## 4. Results

Pair-wise joint log histogram (Figure 3) of the DWI derived scalar images revealed distinct correlation patterns in some pairs of images. FA–AP and FA–GFA showed distinct diagonal correlations patterns with each other; while B0 showed no clear pattern of correlation with other image types, with the exception of MDWI. The scalar images FA, GFA and AP are parameter maps derived from diffusion models, whereas B0 and MDWI are directly derived from the DWI image sequence. We thus grouped FA, AP, and GFA based on their intensity histogram correlations as the anisotropic scalar group (ASG), while B0 and MDWI form a directly derived scalar group (DSG).

**Figure 3:**
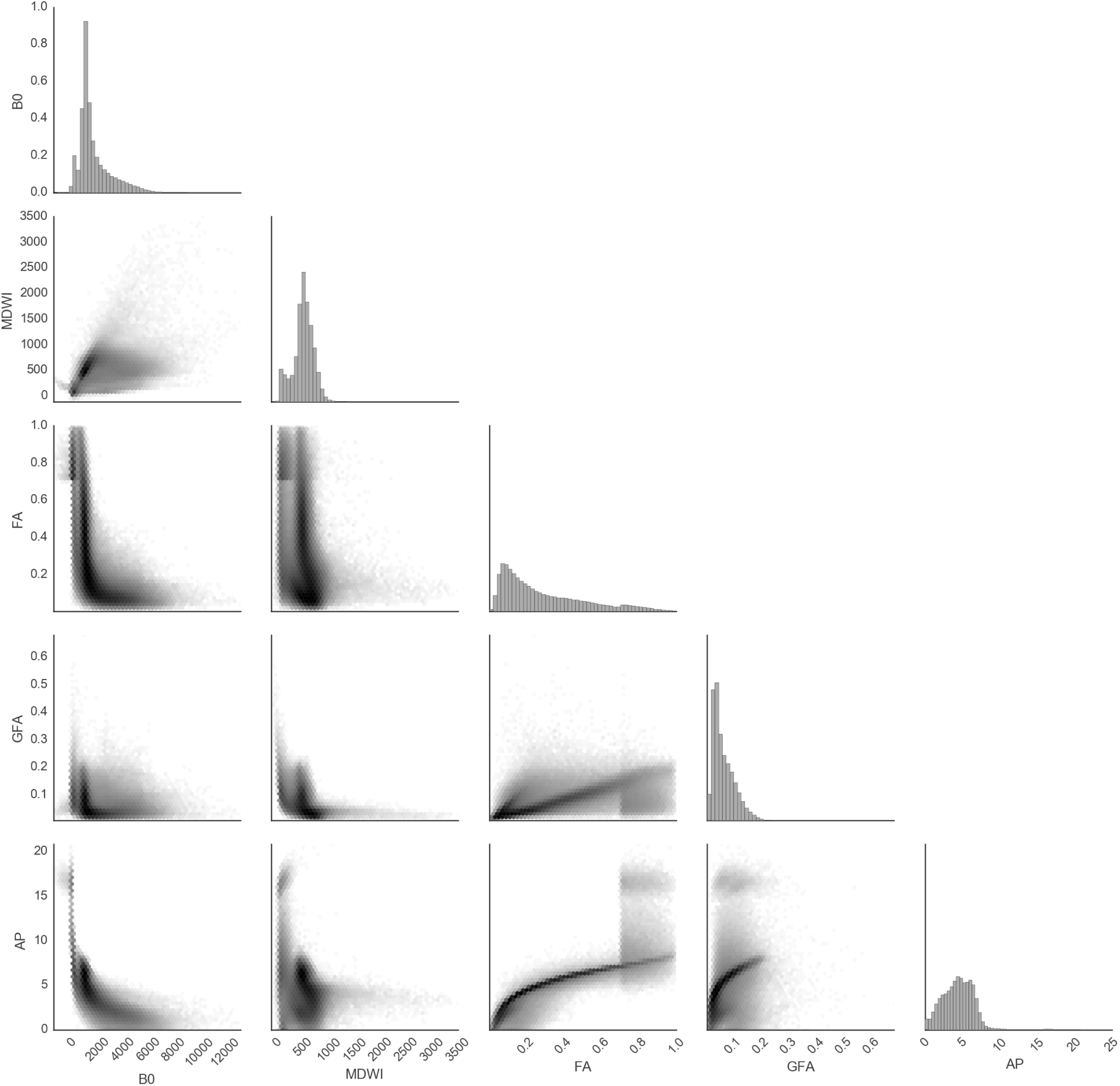
Pair-wise joint log histogram of the DWI derived scalar images.

### 4.1. Differences in initial similarity scores affect registration accuracy

Visual overlay of MDWI to T1 co-registrations (Figure 4) suggests that DWI derived scalar image with higher initial CC and MI scores (Figures 5) appear to result in more accurate registrations in the brainstem, insula, temporal cortices, and lateral ventricles (especially around the ventricular-caudate boundary where CSF/grey/white-matter are found in close proximity) compared to the other image types. Visually, B0, FA and GFA show registration inaccuracies fitting the areas surrounding the brainstem and the lateral ventricles (Figure 4), while MDWI and AP result in improved registration in these key areas. The results of MDWI and AP are visually very similar.

**Figure 4:**
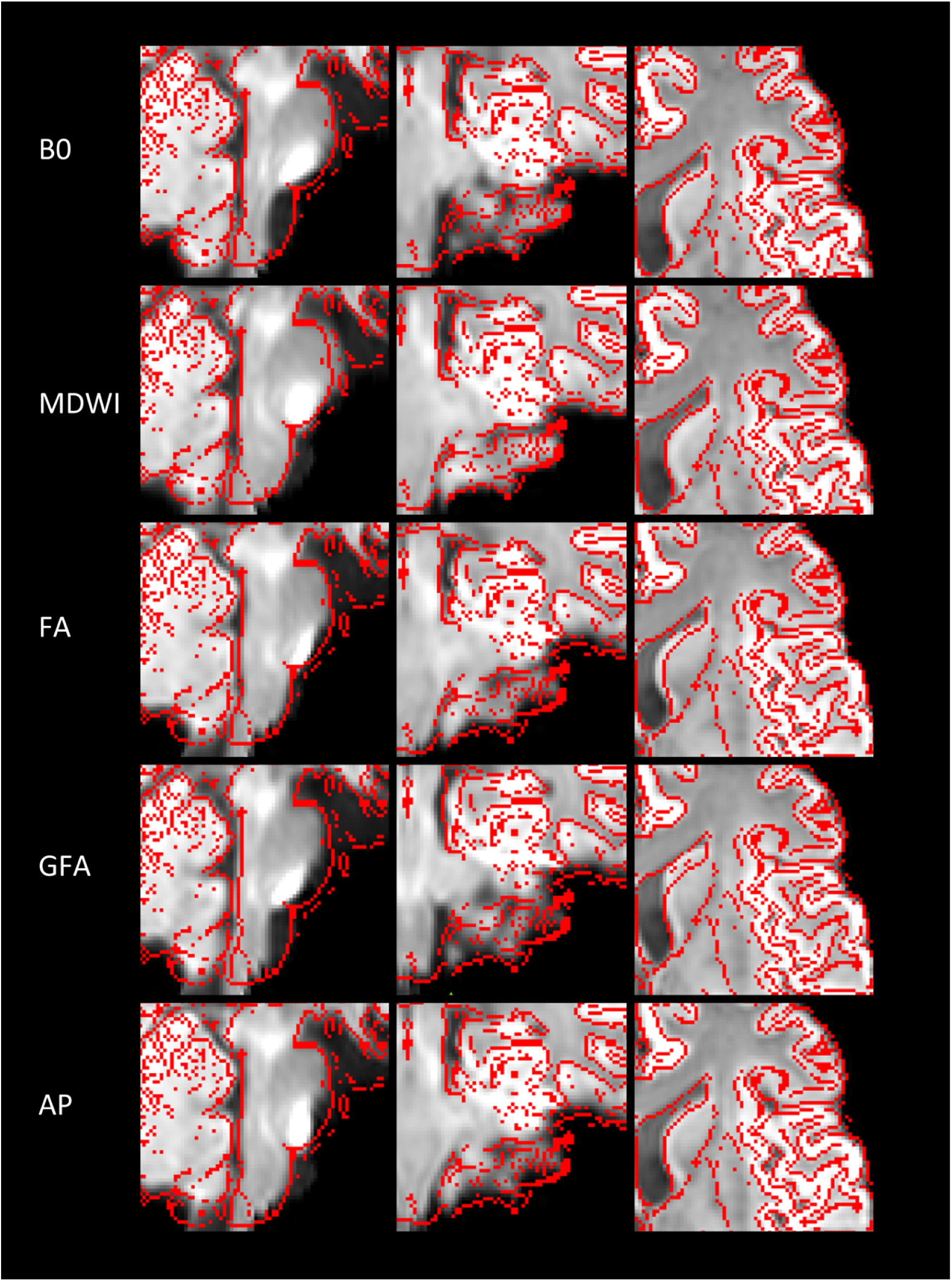
Visual comparisons of MDWI to T1 co-registrations. Shown are examples of MDWI images from one subject deformed using different registration transforms derived from the different image intermediates. T1 is overlaid as thresholded outlines in red. Primary differences in registration quality can be found in the brainstem (left and middle columns), and in the lateral ventricles, especially in the ventricular-caudate boundary (right column).

**Figure 5:**
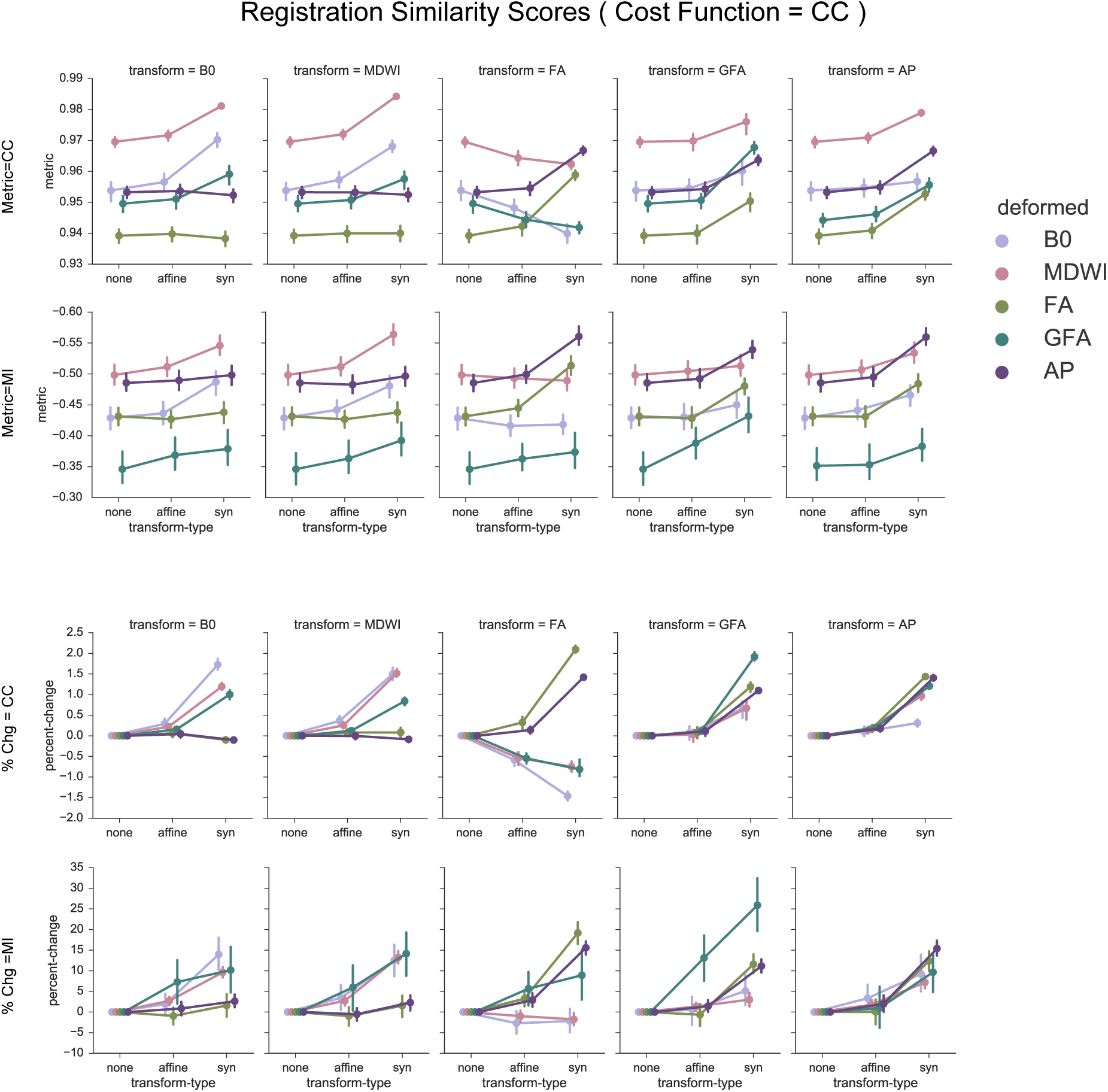
Similarity scores between deformed dMRI scalar images and T1 following registrations with CC as cost function for SyN. Three steps of the registration process are plotted: Initial identity transformation (none), affine transformation (affine) and affine+SyN transformation (SyN). Top panel shows similarity scores as measured by CC and MI, and bottom panel shows average percent change progressions of CC and MI across dMRI scalar images from T_Affine_ to T_Identity_, and from T_Affine_+SyN to T_Affine_. The error bars denote standard deviation across subjects.

The initial T1-intermediate CC and MI similarity scores without any applied transformation (*T*_identity_) showed varied initial values. MDWI showed the highest initial CC scores, with FA images showing lowest initial CC (Figure 5); AP showed similar initial CC score with B0 and GFA. For MI scores (it is important to note that a more negative MI score implies higher similarity in this study), AP showed very similar initial MI score when compared to MDWI (best), while GFA resulted in the highest (worst) initial MI score.

### 4.2. Registration improvements are not uniformly transferred when applied to other image types

The progression of similarity scores following T_Affine_ and T_Affine_+SyN for each of the scalar images (B0, ADWI, MDWI, FA, GFA, and AP) showed similar trends between CC and MI cost-functions (Figures 5, Figure 7; top rows). In general, the similarities of the images between T1 and transformed DWI scalar images increased marginally after T_Affine_, and substantially after T_Affine_+SyN for most of the co-registration transforms. An exception is FA transforms, where CC scores decreased when the transform was applied to other images (Figure 5; rows 1-2, column 4), with the exception of AP.

**Figure 7:**
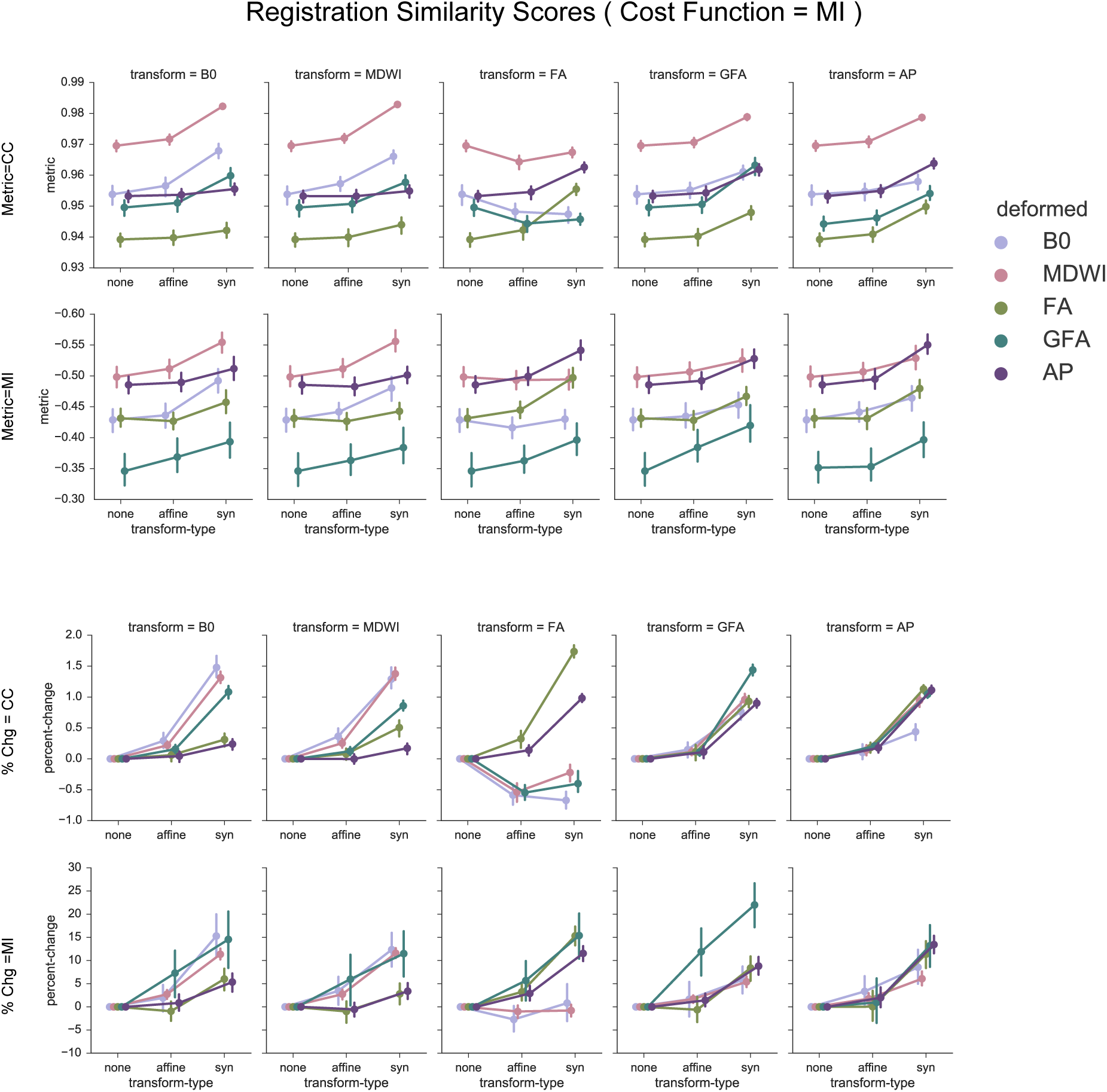
Similarity scores between deformed dMRI scalar images and T1 following registrations with MI as cost function for SyN. Three steps of the registration process are plotted: Initial identity transformation (none), affine transformation (affine) and affine+SyN transformation (SyN). Top panel shows similarity scores as measured by CC and MI, and bottom panel shows average percent change progressions of CC and MI across dMRI scalar images from *T*_Affine_ to *T*_Identity_, and from *T*_Affine_+SyN to *T*_Affine_.

Improvement in CC and MI measures were not uniform for all DWI derived scalar images; this was better illustrated by percent changes (Figures 5; bottom rows). Different intermediate transforms differentially affected the scores in accordance with the degree of intensity correlations as measured in Figure 3. Therefore, DSG transforms preferentially improved DSG, and ASG transforms improved ASG images. FA transforms, however, are notable, where CC scores across all DWI derived scalar images diverged for both CC and MI cost functions (Figures 5, Figure 8; column 4). In fact, FA transforms substantially decreased DSG CC scores, and also negatively impacted DSG MI scores as well. FA transforms also showed exception to GFA, where it affected GFA negatively similar to DSG images. GFA behaved more like DSG images, where B0 and MDWI all favorably affected its CC and MI scores, while at the same time it was strongly affected by AP as well. However GFA seem to behave differently under autodeformation, as evident by its distinct MI changes (Figures 5; column 5). AP transforms showed the least amount of variability when applied to DWI derived scalar images, and consistently improved all the image scores.

**Figure 8:**
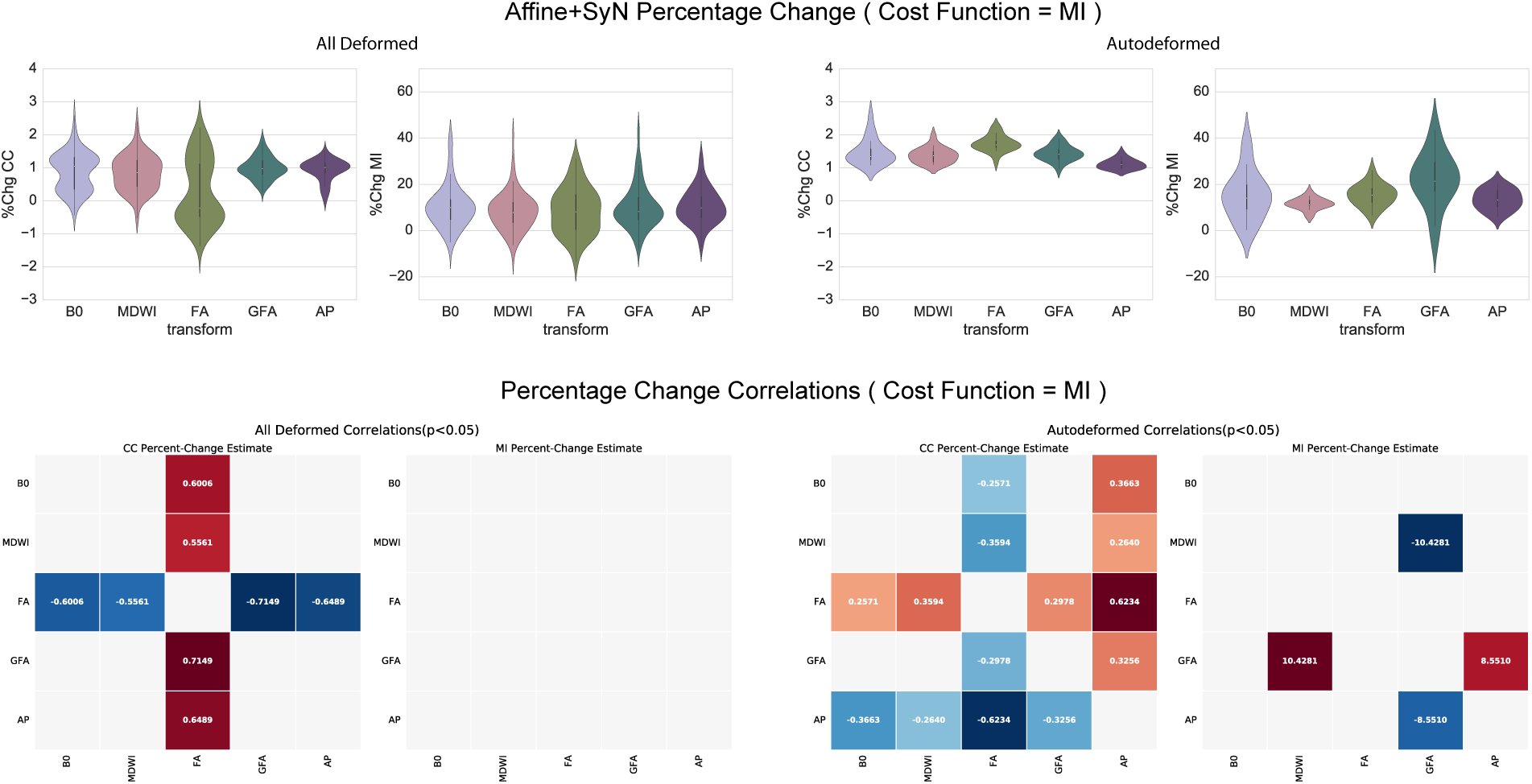
Percent changes of similarity score from *T*_Affine_+SyN to *T*_Identity_ with MI as cost function. Top row shows the violin plot distribution of CC and MI percent change under All Deformations and Autodeformations. Bottom row plots the statistically significant pair-wise correlations of the different transforms. Under all deformations (*D*_all_), CC percent change of FA is shown to be significantly lower (p¡0.05) from that of other transform types. In autodeformations (*D*_auto_). FA has higher percent change. MI percent changes show little differences, and once again highlights the high variability of GFA.

### 4.3. AP-derived transforms show the most consistent improvements when applied to other image types

Between-group statistical comparisons of only T_Affine_+SyN percent change revealed significant findings. Under the CC cost-function (Figure 6), percent change in D_auto_ (Figure 6, bottom panel, column 3) showed that FA and GFA have the highest percent change, while AP has the lowest percent change. With MI scores, D_auto_ showed that only GFA percent changes are significantly greater than others (Figure 6, bottom panel, column 4). GFA, however, also shows the greatest variability in distribution (Figure 6, top panel, column 4). D_all_ in contrast, showed that FA is an outlier where its percentage change is significantly lower than all other images; it is also notable that it shows a distinct bimodal distribution (Figure 6, top panel, column 1); GFA was shown to be significantly higher than B0, MDWI and FA, but not AP. MI percent changes are not significantly different from each other (Figure 6, bottom panel, column 2). MI cost-function results were similar to that of the CC cost-function (Figure 8).

**Figure 6:**
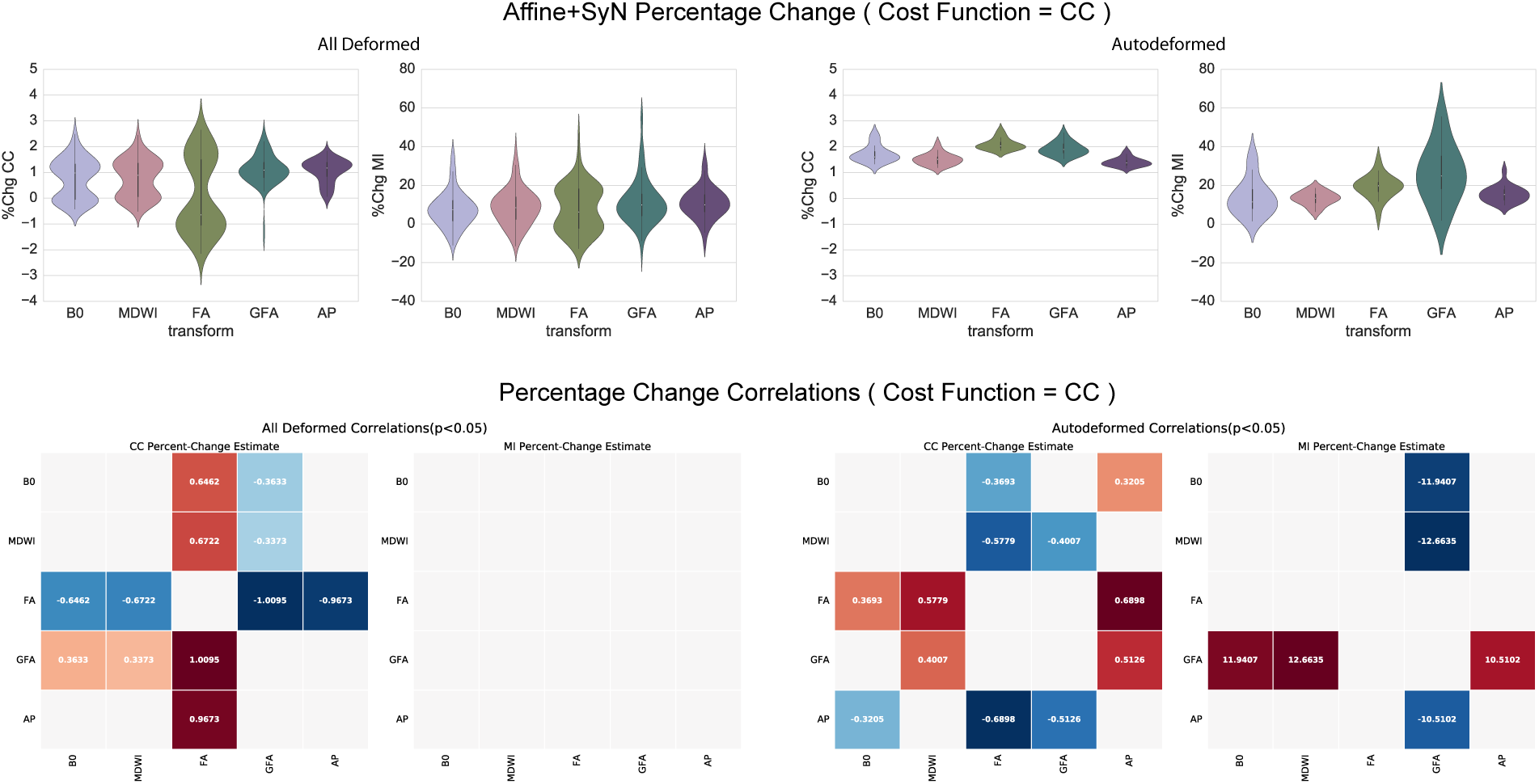
Percent changes of similarity score from T_Affine_+SyN to T_identity_ with CC as cost function. Top row shows the violin plot distribution of CC and MI percent change under All Deformations and Autodeformations. Bottom row plots the statistically significant pair-wise correlations of the different transforms. Under All Deformations (D_all_), CC percent change of FA is shown to be significantly lower (p¡0.05) than that of all other transform types. GFA can be observed to have higher percent change than that of B0, MDWI, and FA. Under Autodeformations (D_auto_), CC percent change of FA and GFA are significantly higher than others, while AP has lower percent change. MI shows little percentage changes between all the images; GFA shows significantly differences under D_auto_, but also shows high variability.

## 5. Discussion

This study presents a comparison of the DWI derived scalar image to use for T1 to DWI co-registration. AP images showed the most consistent improvements in image scores across all of the intermediate scalar images tested, suggesting that AP can offer the most consistent T1-DWI co-registration improvements, while FA and GFA images are poorer choices. We observed, via CC and MI scores, that non-linear registration shows similar trends in registration performance. DWI derived image registration improvements show biases that depend on the intensity correlations with directly derived scalar image group (DSG) or anisotropic scalar image group (ASG) similarities. FA showed significantly lower MI and CC similarity scores that worsened registration results. GFA showed a significant increase in similarity scores, but also greater variability in percent increases when its transform was applied to other image types – suggesting that while this measure is better than FA alone, it is still highly unreliable across subjects.

### 5.1. Registration improvements are biased towards similar image intensity groups

The log joint histogram showed that there is a degree of intensity orthogonality between FA, AP, and GFA as a group (ASG) compared to B0, ADWI and MDWI (DSG), suggesting that there is some connection between co-registration performance and intensity similarity (Figure 3). The greater the intensity similarities between the two images, the greater the effect of the same set of registration transforms. Intensity similarity is not the only factor, however, as evident by the exception to this rule in FA and GFA deformation results (Figure 5). This suggests that intensity orthogonality is not a limiting factor in registration transfer across these scalar images. Therefore, our hypothesis that there exists a better DWI intermediate scalar image that can improve multimodal registration is correct.

### 5.2. Autodeformation is not sufficient to assess registration intermediate performance

Our study demonstrates clear differences in initial similarity scores (i.e. CC and MI) across different DWI scalar images when compared to T1, where MDWI had the highest initial CC and MI scores. Moreover, the AP and MDWI images clearly outperformed the B0, FA, GFA and intermediates in visual inspections.

As expected, autodeformation (e.g. FA to T1 registration applied to the FA image, from greek autos: self) yielded the best metric of similarity compared to when the transform was applied to other image types. The optimal T1-DWI co-registration performance of an intermediate image could be identified by the image’s deformation by its own transform. However, we show that only comparing the autodeformation results is not ideal (Figure 7, 8). Specifically, while the FA autodeformation (using T_Affine_+SyN) performed better compared to all other autodeformations, registrations of other scalar images based on FA transformations clearly resulted in the worst results. Therefore, future comparisons of candidate intermediate images should consider not only the autodeformation, but also the ability of a transform to deform other scalar images. It is also worth noting that affine transformations marginally improve MI scores – which is expected as the head orientations within MR scanners usually do not deviate very far between acquisitions, but most registration improvements stem from corrections to DWI distortions through non-linear registration.

The larger percent change improvements in FA and GFA can be explained by the ceiling effect of registration, where images that are initially more similar to the T1 anatomical image (e.g. MDWI or AP) would have less room for improvements compared to those that are less similar (e.g. FA or GFA). As such, the absolute difference in similarity scores should still be included in the final assessment of registration. It is also important to note that computational algorithms using CC and MI as the optimization metric in the registration process itself cannot fully characterize the goodness-of-fit of one set of images when matched with another. Therefore, our current use of these scores does not preclude the possibility that certain image sub-regions would be better registered using other scalar image types under specific conditions. Moreover, the method of estimating an undistorted DWI with reversed phase-encoding images may still be superior to the use of image intermediates, described here, when this approach is possible. Nonetheless, the post-processing use of intermediate images combined with reverse-phase corrected DWI may result in improved registrations.

As a new DWI derived scalar image, our study is the first to evaluate AP for the purpose of T1-DWI co-registration. AP images can consistently improve similarity outcomes for the various image types and shows more consistent similarity outcomes when compared to other modalities. Further research is needed to determine the optimum method for AP normalization.

### 5.3. Limitations

One important limitation is that the DWI voxel sizes are not isotropic. For this study, the DWI images were acquired at 0.94×0.94×3 mm^3^ voxel resolution. The dataset was acquired on a 3T GE HDx MRI with 8-channels head coil, and therefore does not permit DWI scans of less than 2.6 mm isovoxel resolution at a clinically acceptable scanning time. As such, we compromised on an anisotropic voxel resolution to gain in-plane resolution. Although recent authors have suggested using isotropic up-sampling of MR images to improve image quality (Dyrby et al., 2014), we decided not to introduce up-sampling into our processing pipeline in order to avoid the possibility of additional interpolation-introduced artifacts that may bias towards certain image types. Clinical datasets are often acquired under time and resource constraints, and therefore we believe our findings will be novel for the application of clinical T1-DWI co-registrations at less than ideal conditions.

## 6. Conclusions

This study demonstrated that for T1-DWI co-registration, using AP as the DWI derived scalar image type resulted in improved registration performance. AP offers more consistent registration across DWI derived scalar images. AP requires the calculations of spherical harmonic coefficients as a preprocessing step, and therefore stabilizes when there are greater than 28 gradient directions. It is thereby unsuitable for some legacy DWI datasets acquired with fewer than 28 directions. MDWI can be readily obtained from existing DWI sequences, and is therefore suitable for all existing datasets. We generally recommend the use of AP where available. In the case where calculation of AP images is not possible, MDWI is a viable alternatives as T1-DWI co-registration intermediates. FA and GFA were the poorest performers, and we recommend not using them for registration purposes.

## 7. Acknowledgements

This investigation was supported by a Doctoral Studentship from the Multiple Sclerosis Society of Canada (EGID 2015). This investigation was supported by an Operating Grant from the Multiple Sclerosis Society of Canada (EGID 1712).

## References

Andersson, J. L., & Sotiropoulos, S. N. (2015). An integrated approach to correction for off-resonance effects and subject movement in diffusion MR imaging. NeuroImage, 125, 1063–78.

Andersson, J. L. R., Skare, S., & Ashburner, J. (2003). How to correct susceptibility distortions in spin-echo echo-planar images: application to diffusion tensor imaging. NeuroImage, 20, 870–88.

Avants, B., Tustison, N., & Song, G. (2009). Advanced Normalization Tools (ANTS). Insight Journal, (pp. 1–35).

Avants, B. B., Epstein, C. L., Grossman, M., & Gee, J. C. (2008). Symmetric diffeomorphic image registration with cross-correlation: evaluating automated labeling of elderly and neurodegenerative brain. Medical image analysis, 12, 26–41.

Avants, B. B., Tustison, N. J., Song, G., Cook, P. a., Klein, A., & Gee, J. C. (2011). A reproducible evaluation of ANTs similarity metric performance in brain image registration. NeuroImage, 54, 2033–44.

Basser, P. J., & Jones, D. K. (2002). Diffusion-tensor MRI: theory, experimental design and data analysis - a technical review. NMR in biomedicine, 15, 456–67.

Basser, P. J., & Pierpaoli, C. (2011). Microstructural and physiological features of tissues elucidated by quantitative-diffusion-tensor MRI. 1996. Journal of magnetic resonance (San Diego, Calif.: 1997), 213, 560–70.

Bhushan, C., Haldar, J. P., Choi, S., Joshi, A. A., Shattuck, D. W., & Leahy, R. M. (2015). Co-registration and distortion correction of diffusion and anatomical images based on inverse contrast normalization. NeuroImage, 115, 269–80.

Brown, L. G. (1992). A survey of image registration techniques. ACM Computing Surveys, 24, 325–76.

Cao, Q., Shu, N., An, L., Wang, P., Sun, L., Xia, M.-R., Wang, J.-H., Gong, G.-L., Zang, Y.-F., Wang, Y.-F., & He, Y. (2013). Probabilistic diffusion tractography and graph theory analysis reveal abnormal white matter structural connectivity networks in drug-naive boys with attention deficit/hyperactivity disorder. The Journal of neuroscience: the official journal of the Society for Neuroscience, 33, 10676–87.

Chen, D. Q., DeSouza, D. D., Hayes, D. J., Davis, K. D., O’Connor, P., & Hodaie, M. (2016a). Diffusivity signatures characterize trigeminal neuralgia associated with multiple sclerosis. Multiple sclerosis (Houndmills, Basingstoke, England), 22, 51–63.

Chen, D. Q., Strauss, I., Hayes, D. J., Davis, K. D., & Hodaie, M. (2015). Age-related changes in diffusion tensor imaging metrics of fornix subregions in healthy humans. Stereotactic and Functional Neurosurgery, 93, 151–9.

Chen, D. Q., Zhong, J., Hayes, D. J., Behan, B., Walker, M., Hung, P. S.-P., & Hodaie, M. (2016b). Merged Group Tractography Evaluation with Selective Automated Group Integrated Tractography. Frontiers in neuroanatomy, 10, 96.

Ciccarelli, O., Catani, M., Johansen-Berg, H., Clark, C., & Thompson, A. (2008). Diffusion-based tractography in neurological disorders: concepts, applications, and future developments. Lancet Neurol., 7, 715–27.

Crum, W. R. (2004). Non-rigid image registration: theory and practice.

Dell’Acqua, F., Lacerda, L., Catani, M., & Simmons, A. (2014). Anisotropic Power Maps: A diffusion contrast to reveal low anisotropy tissues from HARDI data. In Proceedings of International Society for Magnetic Resonance in Medicine. Milan, Italy volume 22.

Descoteaux, M., Angelino, E., Fitzgibbons, S., & Deriche, R. (2006). Apparent diffusion coefficients from high angular resolution diffusion imaging: Estimation and applications. Magnetic Resonance in Medicine, 56, 395–410.

Descoteaux, M., Angelino, E., Fitzgibbons, S., & Deriche, R. (2007). Regularized, fast, and robust analytical Q-ball imaging. Magnetic resonance in medicine: official journal of the Society of Magnetic Resonance in Medicine / Society of Magnetic Resonance in Medicine, 58, 497–510.

Dyrby, T. B., Lundell, H., Burke, M. W., Reislev, N. L., Paulson, O. B., Ptito, M., & Siebner, H. R. (2014). NeuroImage Interpolation of diffusion weighted imaging datasets. NeuroImage, 103, 202–13.

Frank, L. R. (2002). Characterization of anisotropy in high angular resolution diffusion-weighted MRI. Magnetic Resonance in Medicine, 47, 1083–99.

Garyfallidis, E., Brett, M., Amirbekian, B., Rokem, A., van der Walt, S., Descoteaux, M., & Nimmo-Smith, I. (2014). Dipy, a library for the analysis of diffusion MRI data. Frontiers in neuroinformatics, 8, 8.

Gupta, V., Malandain, G., Ayache, N., & Pennec, X. (2016). A Framework for Creating Population Specific Multimodal Brain Atlas Using Clinical T1 and Diffusion Tensor Images. In A. Fuster, A. Ghosh, E. Kaden, Y. Rathi, & M. Reisert (Eds.), Computational Diffusion MRI: MICCAI Workshop, Munich, Germany, October 9th, 2015 Mathematics and Visualization (pp. 99–108). Cham: Springer International Publishing.

Hodaie, M., Chen, D. Q. D., Quan, J., & Laperriere, N. (2012). Tractography delineates microstructural changes in the trigeminal nerve after focal radiosurgery for trigeminal neuralgia. PloS one, 7, e32745.

Hodaie, M., Quan, J., & Chen, D. Q. (2010). In vivo visualization of cranial nerve pathways in humans using diffusion-based tractography. Neurosurgery, 66, 788–95; discussion 795.

Hunter, J. D. (2007). Matplotlib: A 2D graphic environment. Computing in Science & Engineering, 9, 90–5.

Irfanoglu, M. O., Modi, P., Nayak, A., Hutchinson, E. B., Sarlls, J., & Pierpaoli, C. (2015). DR-BUDDI (Diffeomorphic Registration for Blip-Up blip-Down Diffusion Imaging) method for correcting echo planar imaging distortions. NeuroImage, 106, 284–99.

Jenkinson, M., & Smith, S. (2001). A global optimisation method for robust affine registration of brain images. Medical Image Analysis, 5, 143–56.

Klein, A., Andersson, J., Ardekani, B. a., Ashburner, J., Avants, B., Chiang, M. C., Christensen, G. E., Collins, D. L., Gee, J., Hellier, P., Song, J. H., Jenkinson, M., Lepage, C., Rueckert, D., Thompson, P., Vercauteren, T., Woods, R. P., Mann, J. J., & Parsey, R. V. (2009). Evaluation of 14 nonlinear deformation algorithms applied to human brain MRI registration. NeuroImage, 46, 786–802.

Leemans, A., & Jones, D. K. (2009). The B-matrix must be rotated when correcting for subject motion in DTI data. Magnetic Resonance in Medicine, 61, 1336–49.

Lenglet, C., Campbell, J. S. W., Descoteaux, M., Haro, G., Savadjiev, P., Wassermann, D., Anwander, A., Deriche, R., Pike, G. B., Sapiro, G., Siddiqi, K., & Thompson, P. M. (2009). Mathematical methods for diffusion MRI processing. NeuroImage, 45, S111–22.

Liu, B., Bai, X., Zhou, F., Han, H., & Hou, C. (2013). Mutual information based three-dimensional registration of rat brain magnetic resonance imaging time-series q. Computers and Electrical Engineering, 39, 1473–84.

Maintz, J. B. A., & Viergever, M. A. (1998). A survey of medical image registration. Medical Image Analysis, 2, 1–36.

Malinsky, M., Peter, R., Hodneland, E., Lundervold, A. J., Lundervold, A., & Jan, J. (2013). Registration of FA and T1-Weighted MRI data of healthy human brain based on template matching and normalized cross-correlation. Journal of Digital Imaging, 26, 774–85.

Mansour, A. R., Baliki, M. N., Huang, L., Torbey, S., Herrmann, K. M., Schnitzer, T. J., & Apkarian, a. V. (2013). Brain white matter structural properties predict transition to chronic pain. Pain, 154, 2160–8.

Mattes, D., Haynor, D. R., Vesselle, H., Lewellen, T. K., & Eubank, W. (2003). PET-CT image registration in the chest using free-form deformations. IEEE Trans.Med.Imaging, 22, 120–8.

McGrath, J., Johnson, K., O’Hanlon, E., Garavan, H., Gallagher, L., & Leemans, A. (2013). White Matter and Visuospatial Processing in Autism: A Constrained Spherical Deconvolution Tractography Study. Autism research: official journal of the International Society for Autism Research, (pp. 1–13).

Mercier, L., Fonov, V., Haegelen, C., Del Maestro, H. F., Petrecca, K., & Collins, D. L. (2012). Comparing two approaches to rigid registration of three-dimensional ultrasound and magnetic resonance images or neurosurgery. International Journal of Computer Assisted Radiology and Surgery, 7, 125–36.

Moayedi, M., & Davis, K. D. (2012). Theories of pain: from Specificity to Gate Control. Journal of neurophysiology, (pp. 5–12).

Peng, H., Orlichenko, A., Dawe, R. J., Agam, G., Zhang, S., & Arfanakis, K. (2009). Development of a human brain diffusion tensor template. NeuroImage, 46, 967–80.

Pieper, S., Lorensen, B., Schroeder, W., & Kikinis, R. (2006). The NA-MIC Kit: ITK, VTK, Pipelines, Grids and 3D Slicer as An Open Platform for the Medical Image Computing Community. In 3rd IEEE International Symposium on Biomedical Imaging: Macro to Nano, 2006. (pp. 698–701). IEEE.

Pluim, J. P. W., Maintz, J. B. a., & Viergever, M. a. (2003). Mutual information based registration of medical images: a survey. IEEE Transactions on medical imaging, XX, 1–21.

Rohde, G. K., Barnett, a. S., Basser, P. J., Marenco, S., & Pierpaoli, C. (2004). Comprehensive Approach for Correction of Motion and Distortion in Diffusion-Weighted MRI. Magnetic Resonance in Medicine, 51, 103–14.

Salomons, T. V., Moayedi, M., Weissman-Fogel, I., Goldberg, M. B., Freeman, B. V., Tenenbaum, H. C., & Davis, K. D. (2012). Perceived helplessness is associated with individual differences in the central motor output system. The European journal of neuroscience, 35, 1481–7.

Sboto-Frankenstein, U. N., Lazar, T., Bolster, R. B., Thind, S., Dreessen de Gervai, P., Gruwel, M. L., Smith, S. D., & Tomanek, B. (2013). Symmetry of the fornix using diffusion tensor imaging. Journal of Magnetic Resonance Imaging, 00, n/a–/=/.

Smith, S. M. (2002). Fast robust automated brain extraction. Human Brain Mapping, 17, 143–55.

Tuch, D. S. (2004). Q-ball imaging. Magnetic Resonance in Medicine, 52, 1358–72.

Tustison, N. J., Cook, P. a., Klein, A., Song, G., Das, S. R., Duda, J. T., Kandel, B. M., van Strien, N., Stone, J. R., Gee, J. C., & Avants, B. B. (2014). Large-scale evaluation of ANTs and FreeSurfer cortical thickness measurements. NeuroImage, 99, 166–79.

Wiech, K., Jbabdi, S., Lin, C. S., Andersson, J., & Tracey, I. (2014). Differential structural and resting state connectivity between insular subdivisions and other pain-related brain regions. Pain, 155, 2047–55.

Yeatman, J. D., Dougherty, R. F., Myall, N. J., Wandell, B. a., & Feldman, H. M. (2012). Tract profiles of white matter properties: automating fiber-tract quantification. PloS one, 7, e49790.

